# Role of KCC2-Dependent Potassium Efflux in 4-Aminopyridine-Induced Epileptiform Synchronization

**DOI:** 10.1101/171488

**Authors:** Oscar C González, Zahra Shiri, Giri P Krishnan, Timothy L Myers, Sylvain Williams, Massimo Avoli, Maxim Bazhenov

**Author notes:** To whom correspondence should be addressed: Maxim Bazhenov, Department of Medicine, University of California, San Diego, La Jolla, CA 92093.

## Abstract

A balance between excitation and inhibition is required to maintain stable brain network dynamics. Traditionally, seizure activity is believed to arise from the breakdown of this delicate balance in favor of excitation with loss of inhibition. Surprisingly, recent experimental evidence suggests that this conventional view may be untrue, and that inhibition plays a prominent role in the development of epileptiform synchronization. Here, we explored the role of the co-transporter KCC2 in the onset of inhibitory network-induced seizures. Our experiments in acute mouse brain slices of either sex revealed that optogenetic stimulation of either parvalbumin- or somatostatin-expressing interneurons induced ictal discharges in rodent entorhinal cortex during 4-aminopyridine application. These data point to a proconvulsive role of GABA_A_ receptor signaling that is independent of the inhibitory input location (i.e., dendritic *vs.* somatic). Further, we developed a biophysically realistic network model implementing complex dynamics of the ion concentrations to explore the mechanisms leading to inhibitory network-induced seizures. In agreement with experimental results, we found that stimulation of inhibitory interneurons induced seizure-like activity in a network with reduced potassium A-current. Model predicted that interneuron stimulation triggered interneuron firing that was accompanied by an increase in intracellular chloride and a subsequent KCC2-dependent gradual accumulation of extracellular potassium promoting epileptiform ictal activity. When the KCC2 activity was reduced, stimulation of the interneurons was no longer able to induce ictal events. Overall, our study provides evidence for a proconvulsive role of GABA_A_ receptor signaling that depends on the involvement of the KCC2 co-transporter.

## 1. Introduction

Several studies have recently confirmed that under some conditions inhibitory GABA_A_ receptor signaling may play a prominent role in seizure generation (Lillis et al., 2012; Hamidi and Avoli, 2015; Sessolo et al., 2015; Uva et al., 2015; Yekhlef et al., 2015; Shiri et al., 2016). This evidence is in conflict with the established notion that epileptiform discharges result from excessive glutamatergic signaling due to reduced inhibition (Ben-Ari et al., 1979; Dingledine and Gjerstad, 1980; Schwartzkroin and Prince, 1980). Indeed, inhibitory interneurons discharge action potentials at seizure onset in *in vitro* (Lillis et al., 2012; Uva et al., 2015; Levesque et al., 2016) and *in vivo* experiments (Grasse et al., 2013; Toyoda et al., 2015). Moreover, seizure-like discharges *in vitro* disappear during pharmacological procedures that interfere with GABA_A_ receptor signaling (Avoli et al., 1996; Lopantsev and Avoli, 1998; Uva et al., 2015). In line with this evidence, direct optogenetic activation of inhibitory interneurons during bath application of 4-aminopyridine (4AP) elicits seizure-like discharges *in vitro* (Yekhlef et al., 2015; Shiri et al., 2016). Together, these data suggest that an increase in inhibitory interneuron synchrony leads to the development of paroxysmal, seizure-like activity under conditions of impaired potassium (K^+^) channel activity. However, the mechanisms of this action remain to be fully understood.

Intracellular chloride concentration ([Cl^-^]_i_) has been shown to increase in principal neurons at seizure onset in 4AP treated conditions (Lillis et al., 2012). Such intracellular accumulation of [Cl^-^]_i_, which is presumably due to the aforementioned increase in GABAergic signaling prior to seizure generation, can be accompanied by a large increase in extracellular potassium concentration ([K^+^]_o_) (Krishnan and Bazhenov, 2011). Recently, *in vitro* optogenetic stimulation of inhibitory interneurons was shown to increase [K^+^]_o_ to levels capable of inducing seizure-like discharges (Yekhlef et al., 2015). Elevated levels of [K^+^]_o_ function as a positive feedback loop, increasing overall network excitability and thus may lead to seizure onset and progression (Pedley et al., 1974; Traynelis and Dingledine, 1988; Somjen, 2002; Frohlich and Bazhenov, 2006; Frohlich et al., 2008; Krishnan and Bazhenov, 2011; González et al., 2015). Previous work has shown that oscillations of [K^+^]_o_ mediate periodic transitions between fast runs and spike-and-wave complexes during seizures (Frohlich and Bazhenov, 2006; Frohlich et al., 2008; Krishnan and Bazhenov, 2011), and that increases in baseline [K^+^]_o_ fluctuations may occur in the presence of cortical trauma (González et al., 2015). Additionally, K^+^ dynamics have been implicated in the transition to seizure and spreading depression, two network states previously thought to be mechanistically distinct (Wei et al., 2014).

The potassium-chloride co-transporter isoform 2 (KCC2) has been proposed as the critical link between the increase in [Cl^-^]_i_ and subsequent increase in [K^+^]_o_ (Rivera et al., 2005; Hamidi and Avoli, 2015; Shiri et al., 2016). Indeed, reduction of KCC2 activity has been shown to prevent the generation of seizures induced by 4AP (Hamidi and Avoli, 2015), as well as the increases in [K^+^]_o_ that occur in response to high-frequency stimulation (Viitanen et al., 2010). Therefore, it has been postulated that synchronized GABAergic activity may cause a gradual accumulation of [Cl^-^]_i_, leading to the activation of KCC2. This activation results in the extrusion of both Cl^-^ and K^+^, allowing K^+^ to accumulate to levels necessary to elicit seizures (Avoli and de Curtis, 2011; Avoli et al., 2016).

In this study, we tested this hypothesis by employing a biophysically realistic network model with dynamic ion concentrations, Na+/K^+^ ATPase activity, and KCC2 activity. We found that reduction of the outward K^+^ (type A) current (I_A_), mimicking the effects of 4AP application, shifted the network dynamics to a state that could transition to seizure-like activity in response to interneuron stimulation. Importantly, the reduction of KCC2 activity (*cf*.,(Hamidi and Avoli, 2015) prevented seizure generation, thus supporting our prediction about the role of KCC2 in ictogenesis.

## 2. Methods

### 2.1. Animals

All procedures were performed according to protocols and guidelines of the Canadian Council on Animal Care and were approved by the McGill University Animal Care Committee. PV-Cre (Jackson Laboratory, B6;129P2-*Pvalb^tm1(cre)Arbr^*/J, stock number 008069) and SOM-Cre (Jackson Laboratory, *Ssttm2.1(cre)Zjh*/J, stock number 013044) homozygote mouse colonies were bred and maintained in house in order to generate pups that were used in this study.

### 2.2. Stereotaxic virus injections

Four PV-Cre (2 male and 2 female) and five SOM-Cre (3 male and 2 female) pups were anesthetized at P15 using isoflurane and positioned in a stereotaxic frame (Stoelting). AAVdj-ChETA-eYFP virus (UNC Vector Core) was delivered in the entorhinal cortex (EC) (0.6 μL at a rate of 0.06 μLImin). Injection coordinates were: anteroposterior -4.00 mm from bregma, lateral +/- 3.60 mm, dorsoventral -4.00 mm. The transverse sinus was used as a point of reference, and the injection needle was inserted with a 2° anteroposterior angle. After completion of the surgery, pups were returned to their home cage.

### 2.3. Brain slice preparation

Mice were deeply anesthetized with inhaled isoflurane and decapitated at P30-40. Brains were quickly removed and immersed in ice-cold slicing solution containing (in mM): 25.2 sucrose, 10 glucose, 26 NaHCO**3**, 2.5 KCl, 1.25 KH_2_PO_4_, 4 MgCl_2_, and 0.1 CaCl_2_ (pH 7.3, oxygenated with 95% O_2_/5% CO_2_). Horizontal brain sections (thickness = 400 μm) containing the EC were cut using a vibratome (VT1000S, Leica) and incubated for one hour or more in a slice saver filled with artificial cerebrospinal fluid (ACSF) of the following composition (in mM): 125 NaCl, 25 glucose, 26 NaHCO_3_, 2 KCl, 1.25 NaH_2_PO_4_, 2 MgCl_2_, and 1.2 CaCl_2_.

### 2.4. Electrophysiological recordings, photostimulation, and analysis

Slices were transferred to a submerged chamber where they were continuously perfused with oxygenated ACSF (KCl and CaCl_2_ adjusted to 4.5 and 2 mM, respectively) (30 °C, 10-15 mL/min). Field potentials were recorded using ACSF-filled microelectrodes (1-2 MΩ) positioned in the EC in the presence of 4AP (150 μM). Signals were recorded with a differential AC amplifier (AM systems), filtered online (0.1-500 Hz), digitized with a Digidata 1440a (Molecular Devices) and sampled at 5 kHz using the pClamp software (Molecular Devices).

For ChR2 excitation, blue light (473 nm, intensity 35 mW) was delivered through a custom-made LED system, where the LED (Luxeon) was coupled to a 3 mm wide fiber-optic (Edmund Optics) and was placed above the recording region. For optogenetic stimulation of interneurons, light pulses (1 s duration) were delivered at 0.2 Hz for 30 s with a 150 s interval between trains. All reagents were obtained from Sigma-Aldrich and were bath applied. Ictal duration and interval are expressed as mean ± SEM. Data were compared using the Student’s t-test. Results were considered significant if the p-value was less than 0.05.

### 2.5. Principal neuron and interneuron models

Principal or excitatory neurons (PNs) and inhibitory interneurons (INs) were both modeled as two compartment models as described previously in (Bazhenov et al., 2002, 2004; Houweling et al., 2005; Frohlich et al., 2008; Krishnan and Bazhenov, 2011; Volman et al., 2011b; Volman et al., 2011a; González et al., 2015; Krishnan et al., 2015). The membrane potential dynamics for each compartment were modeled by the following equations:

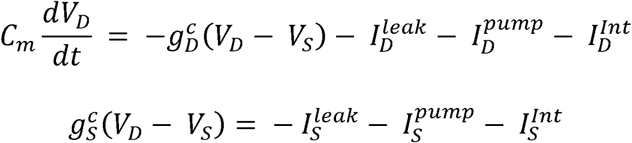

where *V_D_* is the dendritic membrane potential, 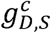 are the dendritic and axosomatic compartment coupling current conductance, 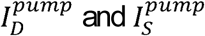 are the sum total Na^+^/K^+^ ATPase currents, 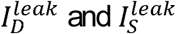 are the sum of the ionic leak currents, and 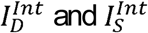 are the intrinsic currents for the dendritic and axosomatic compartments respectively. The intrinsic currents for the dendritic and axosomatic compartments are have been previously described in (Krishnan and Bazhenov, 2011; González et al., 2015; Krishnan et al., 2015).

### 2.6. Dynamic ion concentrations

Ionic concentrations dynamics for [K^+^]_o_, [K^+^]_i_, [Na^+^]_o_, [Na^+^]_i_, [Ca^2^+]_i_, and [Cl^-^]_i_ were modeled similar to our previous work (Krishnan and Bazhenov, 2011; González et al., 2015; Krishnan et al., 2015). In order to model the KCC2 co-transporter, we made some modifications to the [K^+^]_o_ and [Cl^-^]_i_ equations. Briefly, our previous models included KCC2 regulation of [Cl^-^]_i_ in a [K^+^]_o_ dependent manner. However, the [K^+^]_o_ was not affected by KCC2 activity. Our modifications to the [K^+^]_o_ and [Cl^-^]_i_ equations are described as follows:

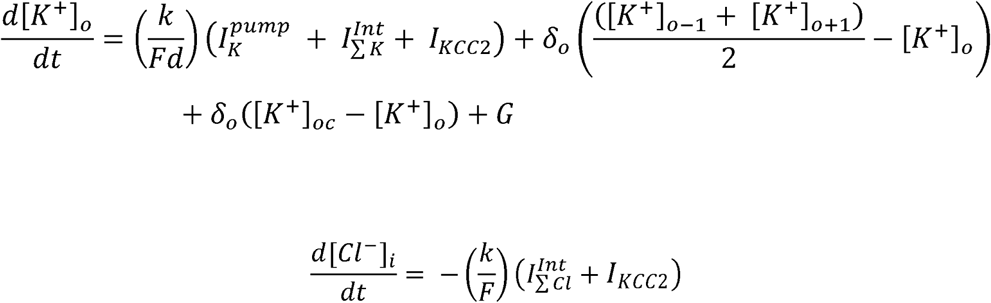

where *F* = 96489 C/mol is the Faraday constant, the ratio of the extracellular volume to surface area is given by *d* = 0.15, and the conversion factor *k* = 10. Additionally, 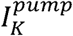 is the K^+^ current through the Na^+^/K^+^ ATPase, [*K*^+^]_*oc*_ is the K^+^ concentration in the adjacent compartment, [*K*^+^]_*o*–1_ and [*K*^+^]_*o*+1_ are the concentrations of K^+^ neighboring cells, and *G* represents the glial buffering of K^+^ as described in detail previously (Krishnan and Bazhenov, 2011; González et al., 2015; Krishnan et al., 2015). Finally, 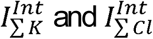 are the sum total intrinsic K^+^ and Cl^-^ currents respectively. *I_KCC2_* defines the efflux of [K^+^]_o_ and [Cl^-^]_i_ generated by the KCC2 co-transporter and is described as follows:

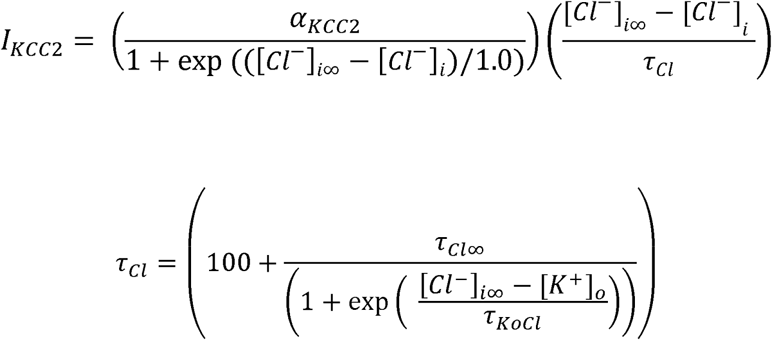

where *α_KCC2_* = 80 defines the strength of the co-transporter, [*Cl*^−^]_*i*∞_ = 5mM is the steady-state Cl^-^ concentration, [*Cl*^−^]_*i*_ is the intracellular Cl^-^ concentration, *τ*_*Cl*∞_ = 4×10^3^s, and *τ_KOCl_* = 0.08s.

### 2.7. Synapse and network properties

We modeled a one dimensional network consisting of 100 PNs and 20 INs. Every PN formed local excitatory synapses onto ten neighboring PNs with AMPA conductance strength of 3.5 nS and NMDA conductance of 0.9 nS. PNs also formed excitatory synaptic connections onto INs with AMPA and NMDA conductance strengths of 2.4 nS and 0.24 nS respectively. INs synapsed onto five local PNs, with GABAA connections of 3.5 nS conductance strength. Additionally, PN and IN received individual afferent excitatory input modeled as a Poisson process as described in our previous studies (Krishnan and Bazhenov, 2011; González et al., 2015; Krishnan et al., 2015).

### 2.8. Estimation of seizure threshold

The seizure threshold was determined using a binary search method that employed an iterative procedure as described in (González et al., 2015). Briefly, at each step of the searching algorithm, the strength of the stimulus would be set to the mean of the upper and lower limits, <*P*>. If this stimulus strength was able to elicit seizure, the upper limit would be set to the current value of <*P*>. If <*P*> was unable to elicit a seizure, the lower limit would take the value of <*P*>. The new stimulus strength, <*P*>, would then be computed based on the updated upper and lower limits. This process continued until the difference between the upper and lower limits was less than 0.1. The threshold was determined to be the average of these final limits.

## 3. Results

### 3.1. Optogenetic stimulation of interneurons triggers ictal discharges

Spontaneous 4AP-induced ictal discharges were recorded from the EC of PV-Cre mice that were transcranially injected with the enhanced ChR2 opsin, ChETA (n = 5 slices). These discharges occurred every 158.07 ± 10.30 s with an average duration of 45.97 ± 1.33 s (n = 124 events). Using a 30 s train of 1 s light pulses at 0.2 Hz that optogenetically activated fast-spiking parvalbumin (PV)-positive interneurons, we were able to trigger ictal discharges of similar duration (i.e., 43.91 ± 2.27 s) but more frequently, at an average interval of 139.45 ± 4.79 s (n = 35 events; *p* < 0.05; figure 1A).

**Figure 1.**
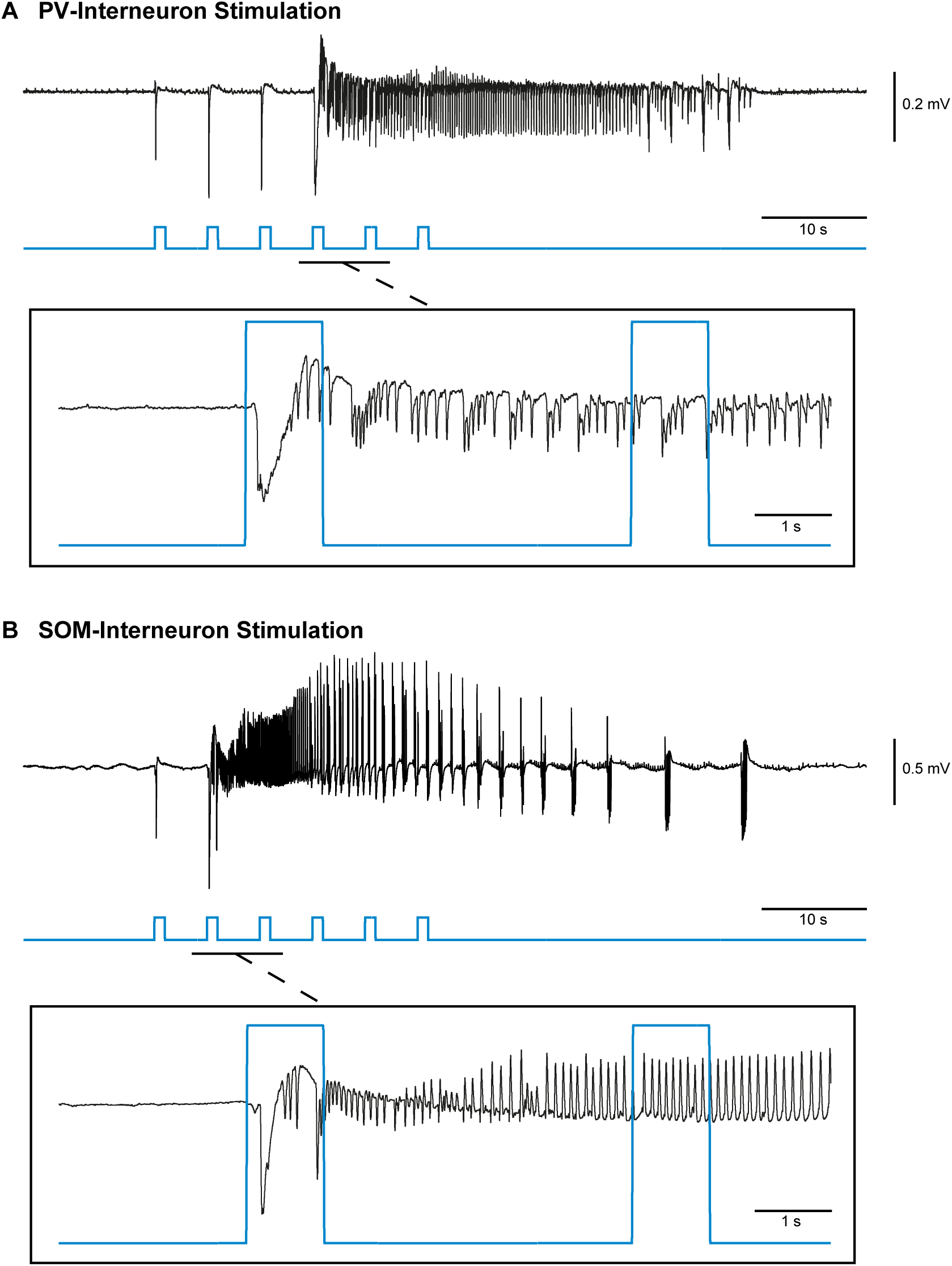
Ictal discharges can be triggered by optogenetic stimulation of PV- or SOM-positive interneurons. **A**, Ictal discharge evoked by a 0.2 Hz series of 1 s light pulses stimulating PV-positive interneurons during bath application of 4AP; ictal onset is expanded to show the timing of the light pulse in relation to ictal onset (box). **B**, The same stimulation parameters applied to SOM-positive interneurons also triggers ictal discharges.

Next, we established whether the ability of interneuron activation to drive ictal discharges was linked exclusively to fast-spiking PV-positive interneurons, or whether ictal discharges could also be triggered by activating regular-spiking somatostatin (SOM)-positive interneurons. Therefore, we obtained brain slices containing the EC of Som-Cre mice that had been transcranially injected with the ChETA opsin (n = 8 slices). Spontaneous 4AP-induced ictal discharges in these experiments occurred every 127.35 ± 6.30 s and lasted on average 63.28 ± 1.84 s (n = 126 events). Using the same protocol used to stimulate PV-interneurons (i.e. 1 s light pulses at 0.2 Hz for 30 s), we were able to trigger ictal discharge of similar duration (55.71 ± 1.90 s) but at a shorter interval of 104.56 ± 5.47 s (n = 44 events; *p* < 0.05; figure 1B). The ictal discharges elicited by the optogenetic activation of either PV- or SOM-expressing interneurons showed characteristic properties of low-voltage, fast (LVF) ictal discharges. Previously, we showed that these LVF ictal discharges are different from the hypersynchronous (HYP) ictal discharges induced by optogenetic stimulation of CamKII-positive principal neurons suggesting a different mechanism between principal neuron- and inhibitory interneuron-induced ictal discharges (Shiri et al., 2016).

### 3.2. Reduction of I_A_ primes the network for interneuron-induced seizure

In order to establish the mechanism by which PV- and SOM-interneuron stimulation cause ictal discharges in the EC of brain slices treated with 4AP, we developed a biophysically realistic network model with dynamic ion concentrations, the electrogenic Na^+^/K^+^ pump, and the KCC2 co-transporter. The network contained synaptically coupled principal (excitatory) neurons (PN) and inhibitory interneurons (IN), and the extracellular compartments of these two neuron types were ionically coupled (see Methods). In order to make comparisons between our model and experimental data, we modeled the application of 4AP as resulting in a 50 percent reduction of the outward K^+^ A-current. Additionally, optogenetic stimulation of interneurons was modeled as a series of 1 s pulses at 0.2 Hz for 30 s similar to those used in our *in vitro* experiments; these “activating” pulses were targeted to all interneurons.

**Figure 2.**
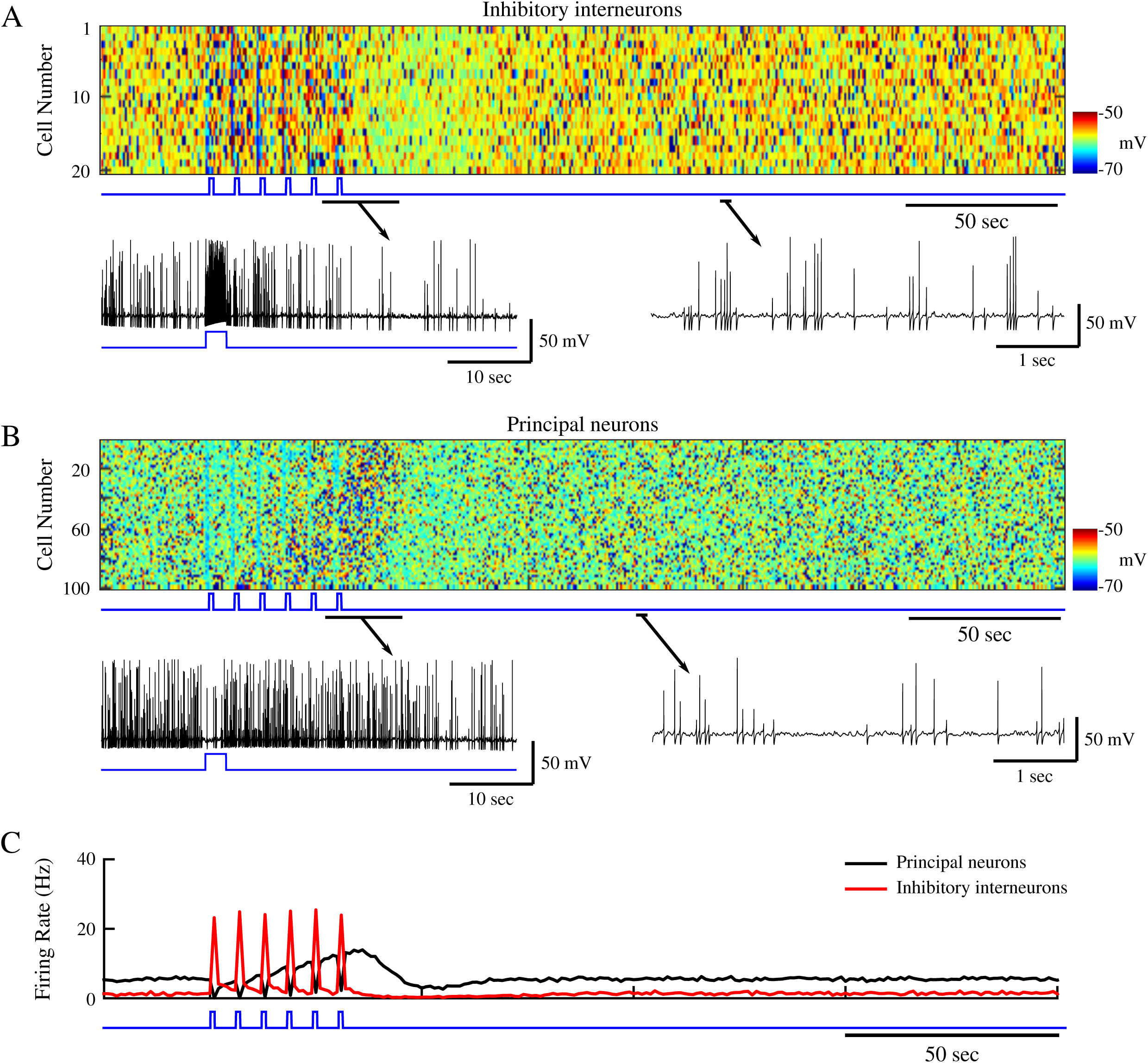
Stimulation of inhibitory interneurons in a healthy network results in brief silencing of excitatory neurons. **A**, Top panel shows raster plot of interneuron activity. Bottom panel shows zoom in of single interneuron spiking from the network in the top panel. Time of interneuron stimulation is indicated by the blue trace. **B**, Principal neuron network activity with zoom in the spiking pattern of a single principal neuron. The blue trace indicates the time of interneuron stimulation. **C**, Mean firing rates for the interneurons (red) and principal neurons (black).

In a control network, one without reduction of I_A_, the stimulus pulses to interneurons resulted in increased firing of INs for the duration of the stimulation, and a gradual decay back to baseline firing rates (figure 2A). During baseline, the mean IN firing rate fluctuated around 1 Hz, but during stimulation it reached ~25 Hz (figure 2C, red). The increase in IN firing during stimulus pulses was accompanied by a hyperpolarization and relative silencing of postsynaptic PNs (figure 2B and C, black trace). Similar to the INs, the PNs displayed a gradual return to a baseline mean firing rate of about 5 Hz (figure 2C, black trace). This control network behaved as expected, i.e., the transient increase in IN firing caused a transient hyperpolarization of the PNs followed by a gradual return to baseline activity.

We next tested the effects induced by IN stimulation on the network dynamics during conditions mimicking 4AP application, i.e., a reduction of I_A_ in both PNs and INs that resulted in a slight increase of the mean intrinsic baseline firing rates of these cell types (~ 4 Hz and ~ 12 Hz for INs and PNs respectively; figure 3D). In this conditions, a single stimulus pulse applied to INs produced an expected hyperpolarization and silencing of PN activity. We then proceeded to apply a sequence of stimuli to all INs to model the effect of optogenetic stimulation similar to our experiment in the control network (figure 2). IN firing peaked at ~ 35 Hz during each stimulus pulse (figure 3A and D, red trace), and during the first 2 pulses PNs were hyperpolarized and silenced by the IN-mediated inhibition (figure 3B and C). However, during subsequent stimulation pulses, the mean firing rate of the PNs began to increase (figure 3D, black trace), and the network developed a seizure-like state, which initiated as focal tonic firing (figure 3B, cells 60-100) before spreading to the remainder of the network and eventually transitioning to a clonic bursting phase (figure 3B and C). Seizure termination was followed by a brief postictal depression and then by the return to baseline firing in both neuron types.

**Figure 3.**
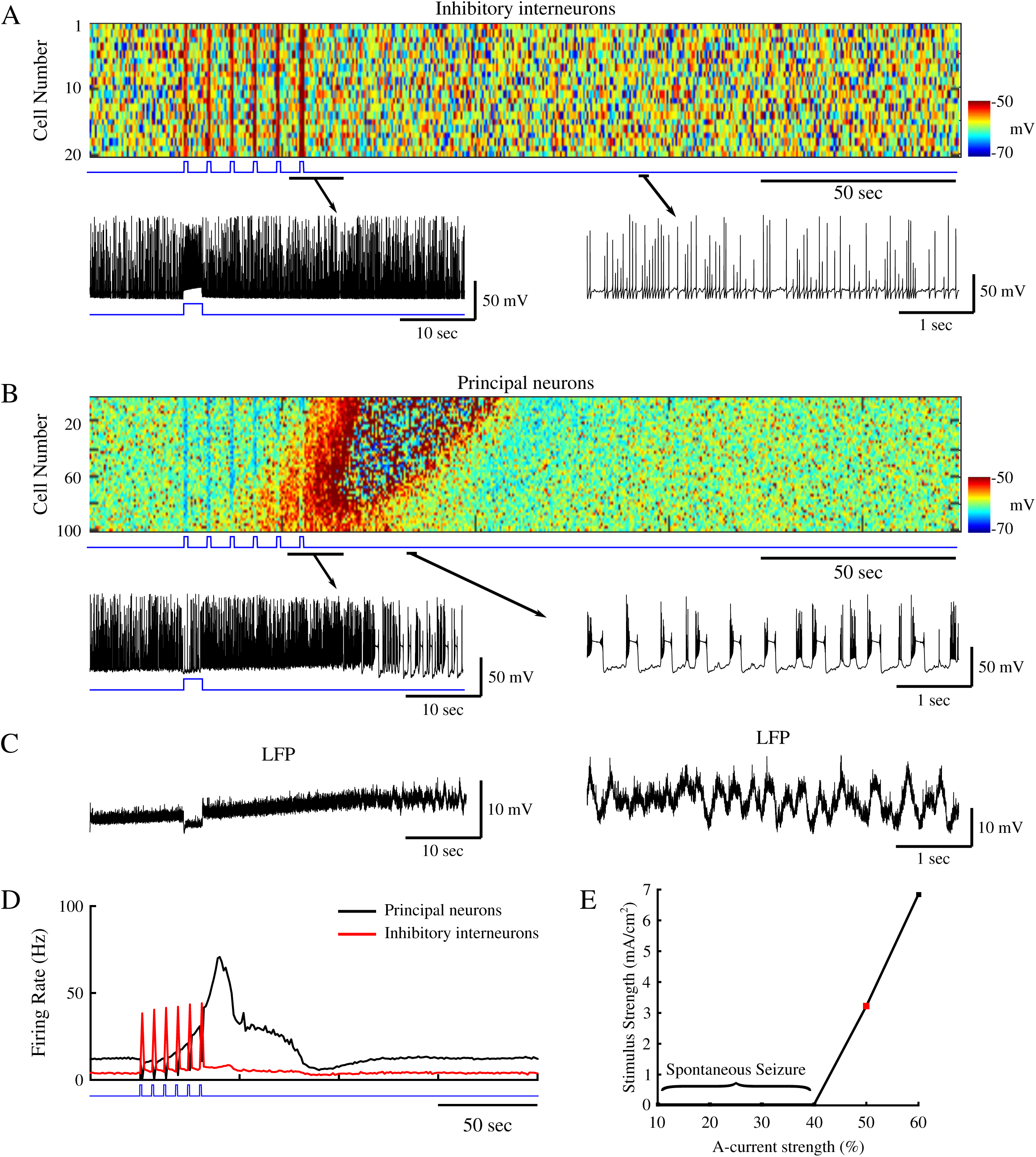
Reduction of A-current increases network excitability allowing for ictogenesis upon interneuron stimulation. **A**, Top panel shows raster plot of interneuron activity. Bottom panel shows zoom in of single interneuron spiking from the network in the top panel. Time of interneuron stimulation is indicated by the blue trace. **B**, Principal neuron network activity with zoom in the spiking pattern of a single principal neuron. The blue trace indicates the time of interneuron stimulation. **C**, Corresponding local field potentials (LFP) for the zoom-ins in B. **D**, Mean firing rates for the interneurons (red) and principal neurons (black). **E**, Stimulus strength necessary for seizure generation as a function of A-current strength. Red square indicates the A-current strength used for the network presented in panels A-D.

Since reduction of I_A_ shifted the network to a state where synchronous inhibitory activity could induce seizure, we next tested the effect of I_A_ strength on the seizure threshold. Reduction of I_A_ made networks more susceptible to seizure (figure 3E; 40-60% range), which can be attributed to increased intrinsic network excitability due to reduced K^+^-dependent repolarization. I_A_ strengths less than 40% of the baseline resulted in spontaneous seizures, while strengths greater than 60% did not show transitions to seizure-like activity following interneuron stimulation.

Dynamics of the [K^+^]_o_ and [Cl^-^]_i_ under the two network conditions – i.e., control and under reduced I_A_ (figures 2 and 3) - revealed stark differences following IN stimulation. As seen in figure 4A, mean [Cl^-^]_i_ and [K^+^]_o_ for all PNs behaved similarly prior to and during IN stimulation under both control and reduced I_A_ conditions (red and black traces respectively). During IN stimulation, both networks revealed increases in the mean [Cl^-^]_i_, and initial decreases in [K^+^]_o_ (figure 4A). The increase in [Cl^-^]_i_ in PNs was presumably due to activation of postsynaptic GABA_A_ receptors, while the transient (initial) decrease in [K^+^]_o_ could reflect the resulting reduction of PN firing. Following termination of the IN stimulation, the [Cl^-^]_i_ concentration in the control network gradually returned to baseline along with the IN firing rate (figure 2C, red trace and figure 4A, red trace in left panel). PN firing became transiently elevated but then also returned to baseline (Fig. 2C, black trace). In contrast, under conditions of I_A_ reduction, IN firing remained elevated and [Cl^-^]_i_ continued to increase following the end of stimulation (figure 2C, red trace and figure 4A, black trace in left panel). Accumulation of [Cl]_i_ caused activation of the KCC2 co-transporter in PNs. As KCC2 uses K^+^ gradient to remove Cl^-^ (Payne et al., 2003), activation of KCC2 led to accumulation of [K^+^]_o_ (figure 4A, right panel, black trace); this last event increased PNs excitability (already elevated under reduced I_A_ conditions) and triggered a positive feedback loop (Frohlich and Bazhenov, 2006; Frohlich et al., 2008; Krishnan and Bazhenov, 2011), initiating transition to seizure activity (figure 4B and 4C).

**Figure 4.**
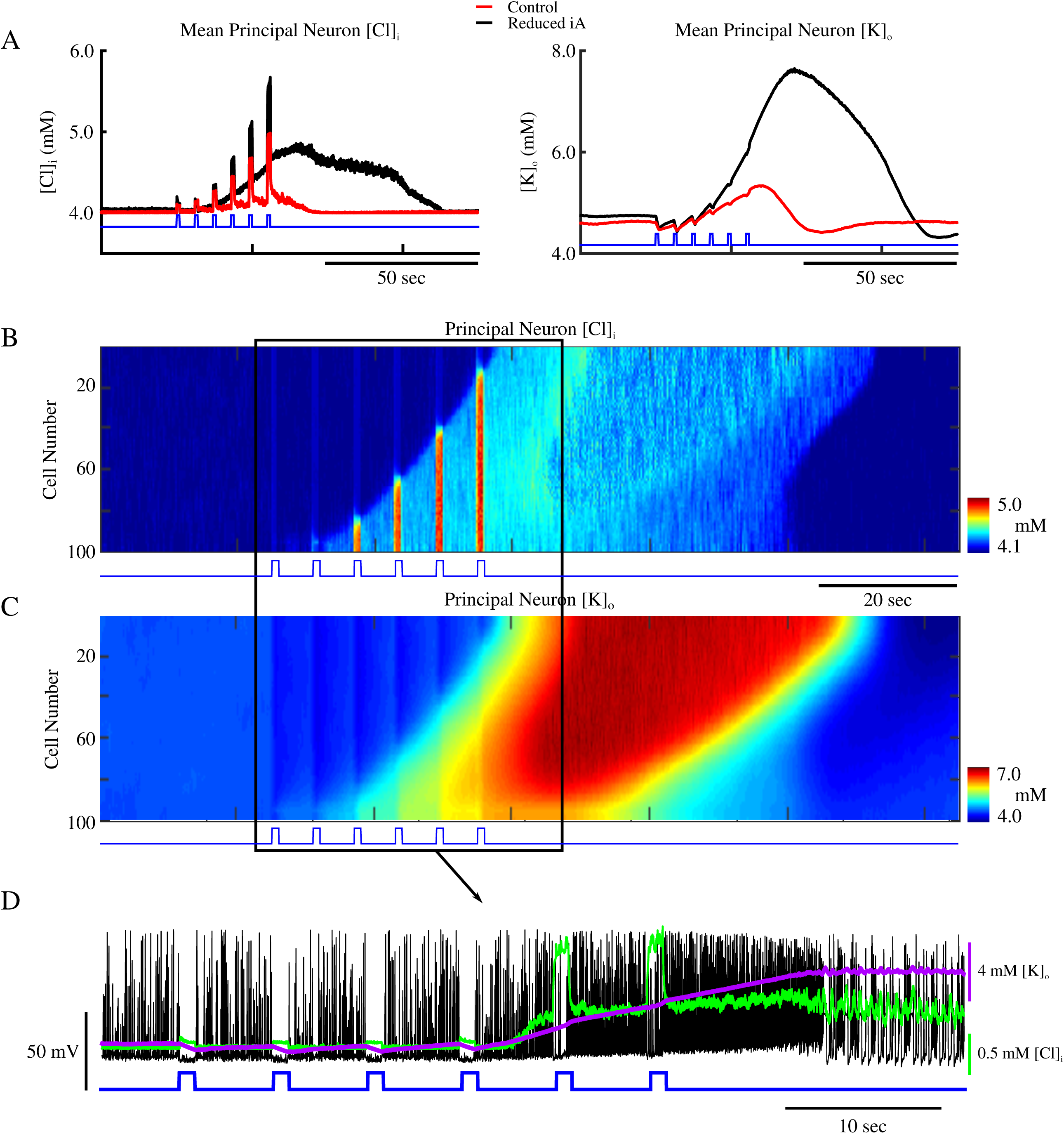
Increased [Cl^-^]_i_ leads to gradual accumulation of [K^+^]_o_ and ictogenesis. **A**, Mean [Cl^-^]_i_ (left) and [K^+^]_o_ (right) for principal neurons in the control network from figure 2, and the reduced A-current network in figure 3 (red and black respectively). The blue trace indicated the onset of interneuron stimulation. **B and C**, network-wide [Cl^-^]_i_ and [K^+^]_o_ for principal neurons. The blue trace indicates the time of interneuron stimulation. **D**, Overlay of the spiking of a single principal neuron (black) from figure 3, and the corresponding [Cl^-^]_i_ (green) and [K^+^]_o_ (purple).

Both [Cl^-^]_i_ in the PNs and [K^+^]_o_ in the surrounding extracellular space remained elevated during the seizure (figure 4B and C, respectively). The expanded sample shown in figure 4D illustrates how activation of interneurons resulted in the silencing and hyperpolarization of PNs during stimulation pulses, while the subsequent accumulation of [K^+^]_o_ increased firing rate and eventually triggered the transition to seizure-like activity. These results suggest that in conditions of increased baseline excitability (such as after reducing I_A_), the network is able to generate seizure-like activity following interneuron activation. Therefore, the mechanism of seizure initiation in our model involved: (a) IN stimulation leading to the release of GABA and postsynaptic activation of GABA_A_ receptors; (b) GABA_A_ activation leading to increase of [Cl^-^]_i_ mediating KCC2 activation; (c) KCC2 activation leading to increase of [K^+^]_o_ sufficiently to initiate the positive feedback loop, thus promoting an “avalanche” increase of excitability.

### 3.3. KCC2 co-transporter activity gives rise to interneuron-induced seizures

To directly test our hypothesis that increased KCC2 activity, resulting from [Cl^-^]_i_ accumulation, may underlie initiation of seizure, we reduced KCC2 co-transporter strength by 50 percent (*α*_*KCC*2_ = 40, see Methods) in a network with reduced I_A_, while stimulating interneurons in a similar way to what was done in the network conditions presented in figure 3. As shown in figure 5A, this resulted in a less excitable network, that was caused presumably by decreased [K^+^]_o_ accumulation due to reduced KCC2 activity level. The IN stimulation was identical to that performed in figures 2 and 3. In this network, stimulation of INs resulted only in the transient silencing of PN activity (figure 5A, top), and [K^+^]_o_ returned to baseline levels shortly after the termination of the IN stimulation (figure 5A, bottom). Unlike the results shown in figure 3, no transition to seizure occurred following termination of the IN stimulation.

**Figure 5.**
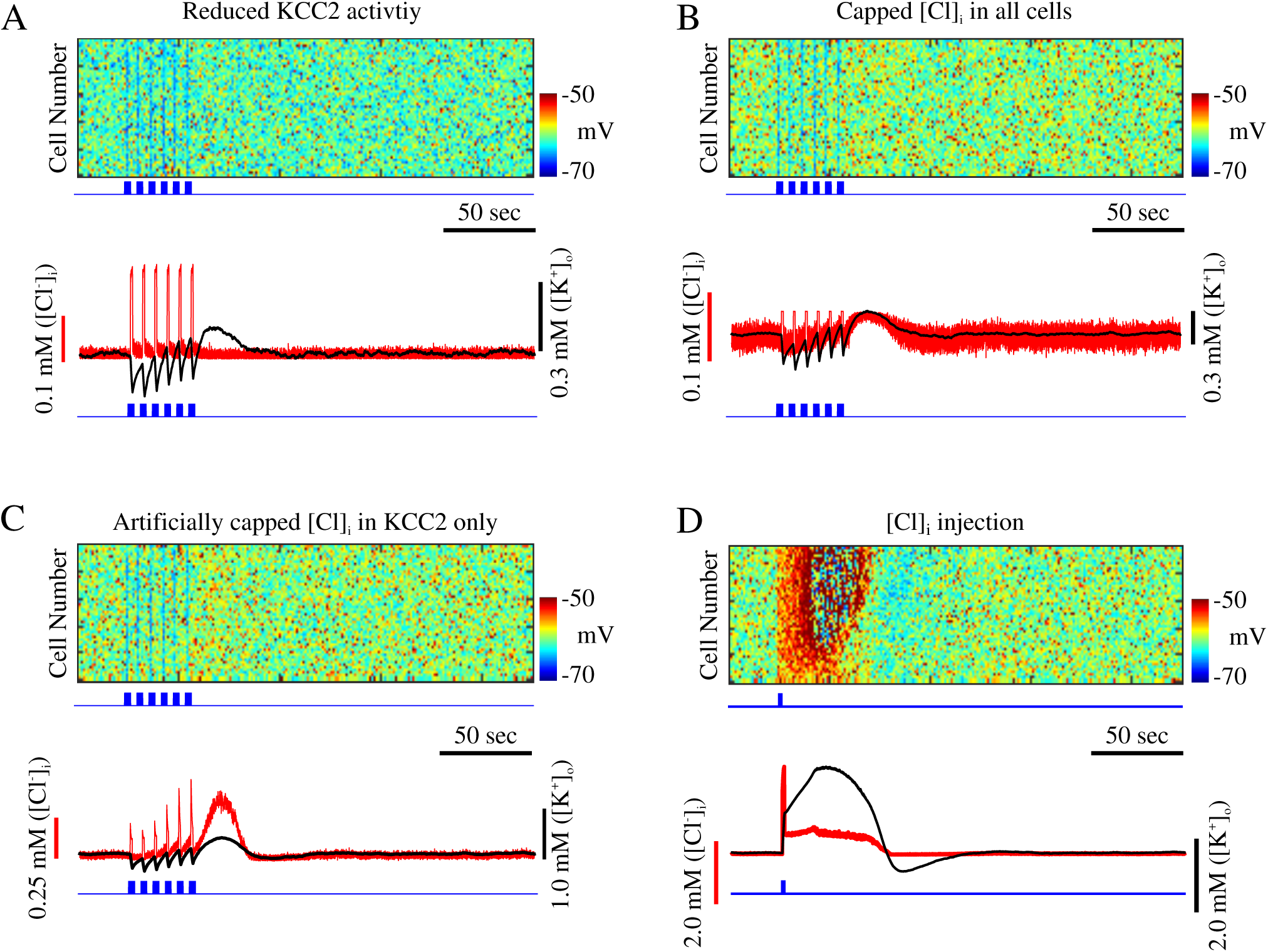
Seizure onset is dependent on KCC2 activation. **A**, Reduction of KCC2 activity prevents seizure. Top, raster plot of principal neurons in a network with reduced A-current and KCC2 activity. Blue trace indicates time of interneurons stimulation. Bottom, corresponding mean [Cl^-^]_i_ (red) and mean [K^+^]_o_ (black) for principal neurons in top panel. **B**, Network with limited [Cl^-^]_i_ accumulation. Top, raster plot showing activity of principal neurons. Bottom, corresponding mean [Cl^-^]_i_ (red) and mean [K^+^]_o_ (black) for principal neurons in top panel. **C**, Network with artificially impaired [Cl^-^]_i_ sensitive K^+^ mechanism. Top, raster plot showing activity of principal neurons. Bottom, corresponding mean [Cl^-^]_i_ (red) and mean [K^+^]_o_ (black) for principal neurons in top panel. **D**, Cl^-^ injection can trigger seizure. Top, rater showing activity of principal neurons. Blue trace shows time of Cl^-^ injection to principal neurons. Bottom, corresponding mean [Cl^-^]_i_ (red) and mean [K^+^]_o_ (black) for principal neurons in top panel.

To further demonstrate that accumulation of [Cl^-^]_i_ was directly responsible for the activation of the KCC2 transporter and subsequent accumulation of [K^+^]_o_, and to compensate for effects of reduction of the baseline activity (Fig. 5A), in the next experiment we kept the strength of KCC2 intact (*α*_*KCC*2_ = 80), but we capped the total amount of [Cl^-^]_i_ that could enter both PNs and INs. In doing so, we limited the maximum amount of [Cl^-^]_i_ that could accumulate in a cell without changing baseline network response or affecting the rate of Cl^-^ influx. In this condition, IN stimulation was also unable to shift the network to a seizure state (figure 5B). [Cl^-^]_i_ increased during the IN stimulation, but since its increase was limited, it was not able to induce a strong enough activation of the KCC2 co-transporter to trigger high [K^+^]_o_ increase. Therefore, following IN stimulation, only a small and brief increase in [K^+^]_o_ was observed (figure 5B, bottom).

Since limiting [Cl^-^]_i_ increase could potentially affect several properties of the model, in the next experiment we artificially limited the effect of [Cl^-^]_i_ on the KCC2 transporter only. Therefore, in this condition, though the [Cl^-^]_i_ could exhibit a significant increase, the KCC2 transporter would only sense a limited increase in [Cl^-^]_i_. That is, the variable in the I_KCC2_ equation representing intracellular Cl^-^ concentration (see Methods) would be less than the actual amount of [Cl^-^]_i_. Essentially, this rendered the K^+^ extrusion mechanism of the KCC2 transporter less sensitive to [Cl^-^]_i_. As shown in figure 5C, IN stimulation was unable to elicit seizure in this network. Brief increases in both [Cl^-^]_i_ and [K^+^]_o_ were observed following the stimulation (figure 5C, bottom). Though the KCC2 activity increased [K^+^]_o_ following IN stimulation, [K^+^]_o_ never reached concentrations sufficient for seizure generation. This last experiment predicts two important points: (a) [Cl^-^]_i_-dependent increase in KCC2-mediated K^+^ extrusion can drive the network to a seizure state, (b) increase of [Cl^-^]_i_ is not sufficient to induce seizure by shifting the GABA_A_ reversal potential.

Finally, we tested whether a direct increase in [Cl^-^]_i_ in PNs could lead to the activation of the KCC2 co-transporter and subsequent elevation of [K^+^]_o_. In this experiment, we used a baseline model where the effect of [Cl^-^]_i_ on KCC2 was intact (similar to network in figure 3). We found that a brief and sufficiently strong increase of [Cl^-^]_i_ in PNs was able to activate KCC2 extrusion of K^+^ thus triggering subsequent seizure generation (figure 5D). Together, these data suggest that the Cl^-^ specific activation of KCC2 activity gives rise to the increase in [K^+^]_o_ sufficient for triggering the transition to seizure.

### 3.4. GABA_A_, and KCC2 influence properties of interneuron-induced seizures

Since our model predicts that GABA_A_ receptor-dependent increase in [Cl^-^]_i_ results in KCC2 mediated increase of [K^+^]_o_ and initiation of seizure, we tested how these two model properties affect seizure onset and duration. By changing the contribution of GABA_A_ to the [Cl^-^]_i_ in the model, we found that decreasing [Cl^-^]_i_ contribution in PNs increased the seizure threshold (figure 6A). Reducing the GABA_A_ receptor-dependent increase in [Cl^-^]_i_ to 94 % of the baseline prevented IN stimulation from inducing seizures in the model. The reduced contribution of GABA_A_ receptor activations to the increase in [Cl^-^]_i_ also resulted in shorter seizure duration (figure 6C). These effects arise from the fact that reduced [Cl^-^]_i_ accumulation leads to lower KCC2 activation and reduced K^+^ efflux.

**Figure 6.**
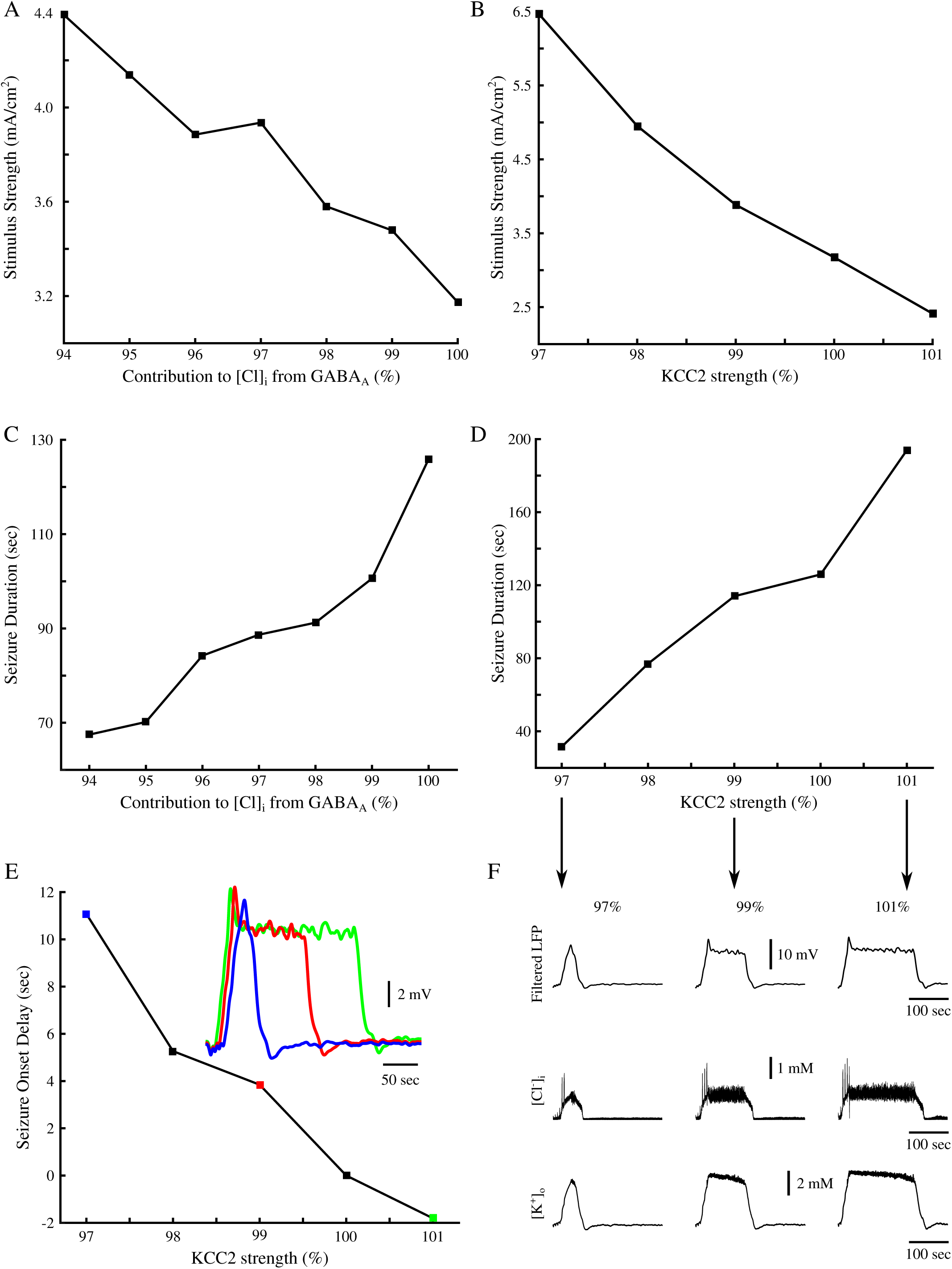
Contribution to [Cl^-^]_i_ from GABA_A_, and KCC2 activity modulate seizure threshold, duration and onset. **A**, Seizure threshold as a function of GABA_A_ contribution to [Cl^-^]_i_. **B**, Seizure threshold as a function of KCC2 strength. **C**, Seizure duration as a function of GABA_A_ contribution to [Cl^-^]_i_. **D**, Seizure duration as a function of KCC2 activity. **E**, Seizure onset delay as a function of KCC2 activity. Delay was measured as the time between the onset times of a seizure in the network with 100% KCC2 strength as compared to the seizure onset times in a network with varied KCC2 strength. Inset shows examples of filtered seizure LFPs. Colored data points correspond to the sampled data in the inset. **F**, Examples of different seizure durations as a result of varied KCC2 strength. Top, filtered network LFP. Middle and Bottom, corresponding mean [Cl^-^]_i_ and [K^+^]_o_ respectively. Arrows point to the corresponding data points in **D**.

This prediction was further validated by directly changing KCC2 strength (*α*_*KCC*2_). As illustrated in figure 6B, increasing KCC2 strength decreased seizure threshold, while decreasing KCC2 strength resulted in an increased threshold. Decreasing KCC2 strength also resulted in decreased seizure durations, while increased KCC2 strength increased seizure duration (figure 6D and F). Interestingly, reduction of KCC2 strength also delayed the onset time to the seizure (figure 6E). In the networks with low KCC2 activity (figure 6F, blue), there was a difference of about 10 sec between seizure onset times in the network with 100 % KCC2 strength and the one with 97 % strength. Accordingly, in the networks with stronger KCC2 (figure 6E, green), seizure onset occurred prior to the seizure onset in a baseline network with 100% KCC2 strength. This effect can be explained by the rate of [K^+^]_o_ accumulation resulting from KCC2 activity. When KCC2 activity was enhanced, [K^+^]_o_ accumulated faster and reached the critical threshold that was sufficient for initiation of seizure after only few seconds.

## 4. Discussion

The K^+^ channel blocker 4AP elicits seizure activity both *in vivo* and *in vitro* (Avoli and de Curtis, 2011). Optogenetic activation of inhibitory interneurons, in acute mouse brain slices exposed to 4AP, can trigger seizure-like discharges (Sessolo et al., 2015; Yekhlef et al., 2015; Shiri et al., 2016). Blocking KCC2 activity with either VU024055 or high doses of bumetanide abolished ictal discharges in 4AP-treated rat brain slices (Hamidi and Avoli, 2015), suggesting that this form of inhibition-induced seizure may involve activation of the KCC2 co-transporter. In this study, we explored the hypothesis that in conditions of elevated cortical excitability (as in the presence of 4AP), Cl^-^-dependent activation of the KCC2 co-transporter makes the network progress to a seizure state by promoting increases in the extracellular K^+^. Our *in vitro* data and computer simulation results suggest that synchronous activation of the inhibitory interneurons can trigger development of paroxysmal activity in the network with reduced potassium A-current.

### 4.1. K^+^ channelopathies in epilepsy

Various channelopathies, including mutated or misregulated K^+^ channels, have been suggested to underlie certain forms of genetic epilepsies (D’Adamo et al., 2013; Lascano et al., 2016). Indeed, mutations in K_V_4 α-subunits are present in some patients suffering from pharmacoresistant temporal lobe epilepsy (Singh et al., 2006; D’Adamo et al., 2013). Specifically, a truncation mutation of the K_V_4.2 α-subunit, responsible for the I_A_, has been observed in a human patient (Singh et al., 2006). This truncation mutation results in an attenuated I_A_ and subsequent increases in seizure susceptibility. In addition to mutations of specific ion channels, mutations of genes encoding proteins which modulate ion channel activity have been found. Patients suffering from autosomal dominant partial epilepsy with auditory features (ADPEAF) have been shown to have point mutations in the leucine-rich glioma-inactivated 1 (LGI1) gene, resulting in reduced neuronal secretion of LGI1 (Ottman et al., 2004; Nobile et al., 2009; Dazzo et al., 2015). Neuronally secreted LGI1 complexes with K_V_1.4 and K_V_β1, two known subunits comprising the A-type channels, prevent rapid inactivation of A-type currents (Schulte et al., 2006). The reduction of LGI1 expression in patients with ADPEAF results in the rapid inactivation of A-type channels and subsequent hyperexcitability.

The I_A_ antagonist, 4AP, has been shown to cause increased neuronal excitability and seizure-like discharges *in vivo* (Fragoso-Veloz et al., 1990; Levesque et al., 2013) and *in vitro* (Avoli et al., 1996; Lopantsev and Avoli, 1998). Interestingly, direct knockout of the K_V_4.2 α-subunit resulted in increased excitability but did not generate spontaneous seizures *in vivo* or *in vitro.* This knockout did, however, increase seizure susceptibility in response to additional proconvulsive pharmacological agents (Barnwell et al., 2009). Previous studies proposed that reduction of A-type K^+^ current promotes ictogenesis by directly increasing neuronal excitability (Galvan et al., 1982; Gustafsson et al., 1982; Yamaguchi and Rogawski, 1992). In contrast, our study predicts that the reduction of A-type K^+^ current leads to increased excitability of both excitatory and inhibitory neurons and that the latter is critical for epileptogenesis. We show that acute brain slices treated with 4AP showed transition to seizure when perturbed by photostimulation of inhibitory interneurons. We hypothesize that the mechanism, by which increased GABAergic signaling may lead to paroxysmal discharges, involves Cl^-^ -dependent activation of KCC2 followed by increases in the extracellular K^+^.

### 4.2. GABA_A_ receptor-dependent [K^+^]_o_ excitatory transients

Early studies proposed that reduced inhibition underlies seizure generation and perhaps epilepsy (Ben-Ari et al., 1979; Dingledine and Gjerstad, 1980; Schwartzkroin and Prince, 1980). Later, this view was challenged in several studies, (de Curtis and Avoli, 2016), which revealed that synchronous inhibitory interneuron activity occurs prior to seizure onset in slices treated with 4AP (Lillis et al., 2012; Uva et al., 2015; Levesque et al., 2016). It has been reported that intense GABAergic stimulation results in increase of [K^+^]_o_ and long-lasting depolarizations (Rivera et al., 2005; Viitanen et al., 2010). Additionally, application of either bicuculline or furosemide inhibits these events (Viitanen et al., 2010). Indeed, these GABAergic excitatory [K^+^]_o_ transients have been shown to elicit prolonged depolarizations in rat CA1 and EC, and may play a prominent role in seizure generation (Lopantsev and Avoli, 1998; Viitanen et al., 2010). The proconvulsive GABAergic excitatory [K^+^]_o_ transients may give rise to the spontaneous seizure onset in patients with K^+^ channel abnormalities.

*In vitro* and *in silico* results presented in this study predict the mechanisms by which GABAergic signaling can trigger seizure onset. We propose that the increased GABAergic signaling, such as triggered by stimulation of inhibitory interneurons, induces Cl^-^ build up, followed by Cl^-^ dependent activation of the KCC2 co-transporter and subsequent increase of [K^+^]_o_. Indeed, high frequency stimulation has been shown to cause increases in [K^+^]_o_ in response to intense GABA_A_ receptor activation (Ruusuvuori et al., 2004; Rivera et al., 2005; Viitanen et al., 2010). Optogenetic stimulation of either SOM- or PV-expressing interneurons also causes large increases in [K^+^]_o_ (Yekhlef et al., 2015). It has been shown that sufficiently large initial increase of [K^+^]_o_ can give rise to a positive feedback loop due to an increase in the network excitability through [K^+^]_o_ dependent depolarization of neurons, which in turn results in a further increase of [K^+^]_o_, which can lead to epileptiform activity (Somjen, 2002; Frohlich and Bazhenov, 2006; Frohlich et al., 2008; Krishnan and Bazhenov, 2011; Wei et al., 2014; González et al., 2015; Krishnan et al., 2015). Out new study predicts that the mechanisms leading to the initial increase in [K^+^]_o_ which kicks the network into the vicious feedback cycle may involve KCC2-dependent efflux of K^+^.

### 4.3. KCC2 in epilepsy

Downregulation of KCC2 expression levels have been suggested to underlie the development of epilepsy in patients (Huberfeld et al., 2007; Buchin et al., 2016). However, other studies have shown that increased KCC2 activation may play a prominent role in seizure generation (Viitanen et al., 2010; Hamidi and Avoli, 2015). Activity dependent regulation of KCC2 expression may explain this seemingly conflicting evidence. Indeed, KCC2 expression has been shown to reduce following increases in activity and epileptiform discharges (Rivera et al., 2002; Rivera et al., 2004; Rivera et al., 2005). Our computational model revealed that the reduction of KCC2 activity prevents seizures in response to intense GABAergic signaling, suggesting that the observed reduction of KCC2 expression may not be a seizure triggering factor but rather a protective mechanism to reduce the likelihood of seizures being triggered by other factors. Our study predicts that increase of [Cl^-^]_i_ in excitatory neurons activates KCC2 co-transporter and promotes initiation of seizure. Consistent with this prediction, recent experimental studies have shown large increases in [Cl^-^]_i_ in excitatory neurons prior to paroxysmal discharges (Lillis et al., 2012). Our model also predicts that increases in KCC2 activity can increase seizure susceptibility and duration. This result is consistent with previous computational modelling and experimental work (Krishnan and Bazhenov, 2011; Hamidi and Avoli, 2015). Importantly, our model predicts a complex effect of GABA_A_ inhibition in seizure development. On one hand, increase of GABA_A_ signaling would act to suppress the network activity, on the other it would promote increase of [Cl^-^]_i_ in excitatory neurons which drives KCC2 activation and [K^+^]_o_ efflux, thus paradoxically increasing network excitability. The balance of these opposite factors determines the resulting network dynamics (normal vs epileptic) in the physiological settings.

## 5. Conclusions

Patients suffering from pharmacoresistant seizures make up approximately 30% of the total number of people living with epilepsy (Nadkarni et al., 2005; Perucca and Tomson, 2011). Among the commonly prescribed antiepileptic drugs, several of them (such as benzodiazepine and barbiturates) enhance GABA_A_ receptor function (Nadkarni et al., 2005; Perucca and Tomson, 2011). Our current study predicts that increased GABAergic signaling, through the treatment with these antiepileptic drugs, may exacerbate existing seizures in patients with K^+^ channelopathies. On the other hand, we present modeling data indicating that modulation of KCC2 activity may prevent seizure generation in these patients. Thus, our findings may provide a new avenue for pharmacological interventions in patients suffering from intractable epilepsy due to K^+^ channelopathies.

## Acknowledgements

This study was supported by NIH grants NS081243, MH099645, EB009282 and Canadian Institutes of Health Research grants 8109 and 74609. Oscar C González is supported by the NSF Graduate Research Fellowship under grant DGE-1326120; Zahra Shiri received a student scholarship from the Savoy Foundation for Epilepsy.

